# Learning long temporal sequences in spiking networks by multiplexing neural oscillations

**DOI:** 10.1101/766758

**Authors:** Philippe Vincent-Lamarre, Matias Calderini, Jean-Philippe Thivierge

**Affiliations:** School of Psychology and Center for Neural Dynamics, University of Ottawa, Ottawa, Ontario, Canada

**Keywords:** Neural oscillations, Spiking neural networks, Recurrent neural networks, Temporal processing, Balanced networks

## Abstract

Many cognitive and behavioral tasks – such as interval timing, spatial navigation, motor control and speech – require the execution of precisely-timed sequences of neural activation that cannot be fully explained by a succession of external stimuli. We show how repeatable and reliable patterns of spatiotemporal activity can be generated in chaotic and noisy spiking recurrent neural networks. We propose a general solution for networks to autonomously produce rich patterns of activity by providing a multi-periodic oscillatory signal as input. We show that the model accurately learns a variety of tasks, including speech generation, motor control and spatial navigation. Further, the model performs temporal rescaling of natural spoken words and exhibits sequential neural activity commonly found in experimental data involving temporal processing. In the context of spatial navigation, the model learns and replays compressed sequences of place cells and captures features of neural activity such as the emergence of ripples and theta phase precession. Together, our findings suggest that combining oscillatory neuronal inputs with different frequencies provides a key mechanism to generate precisely timed sequences of activity in recurrent circuits of the brain.

## 1. Introduction

Virtually every aspect of sensory, cognitive and motor processing in biological organisms involves operations unfolding in time (1). In the brain, neuronal circuits must represent time on a variety of scales, from milliseconds to minutes and longer circadian rhythms (2). Despite increasingly sophisticated models of brain activity, time representation remains a challenging problem in computational modelling (3, 4).

Recurrent neural networks offer a promising avenue to detect and produce precisely timed sequences of activity (5). However, it is challenging to train these networks due to their complexity (6), particularly when operating in a chaotic regime associated with biological neural networks (7, 8).

One avenue to address this issue has been to use reservoir computing (RC) models (9, 10). Under this framework, a recurrent network (the reservoir) projects onto a read-out layer whose synaptic weights are adjusted to produce a desired response. However, while RC can capture some behavioral and cognitive processes (11–13), it often relies on biologically implausible mechanisms, like strong feedback form the readout to the reservoir or implausible learning rules required to control the reservoir’s dynamics. Further, current RC implementations offer little insight to understand how the brain generates activity that does not follow a strict rhythmic pattern (1, 5). That is because RC models are either restricted to learning periodic functions, or require an aperiodic input to generate an aperiodic output, thus leaving the neural origins of aperiodic activity unresolved (5). A solution to this problem is to train the recurrent connections of the reservoir to stabilize innate patterns of activity (12), but this approach is more computationally expensive and is sensitive to structural perturbations (14).

To address these limitations, we propose a biologically plausible spiking recurrent neural network (SRNN) model that receives multiple independent sources of neural oscillations as input. The architecture we propose is similar to previous RC implementations (13), but we use a balanced SRNN following Dale’s law, which are typically used to model the activity of cortical networks (15, 16). The combination of oscillators with different periods creates a multi-periodic code that serves as a time-varying input that can largely exceed the period of any of its individual components. We show that this input can be generated endogenously by distinct sub-networks, alleviating the need to train recurrent connections of the SRNN to generate long segments of aperiodic activity. Thus, multiplexing a set of oscillators into a SRNN provides an efficient and neurophysiologically grounded means of controlling a recurrent circuit (14). Analogous mechanisms have been hypothesized in other contexts including grid cell representations (17) and interval timing (18, 19).

This paper is structured as follows. First, we describe a simplified scenario where a SRNN that receives a collection of input oscillations learns to reproduce an arbitrary time-evolving signal. Second, we extend the model to show how oscillations can be generated intrinsically by oscillatory networks that can be either embedded or external to the main SRNN. Third, we show that a network can learn several tasks in parallel by “tagging” each task to a particular phase configuration of the oscillatory inputs. Fourth, we show that the activity of the SRNN captures temporal rescaling and selectivity, two features of neural activity reported during behavioral tasks. Fifth, we train the model to reproduce natural speech at different speeds when cued by input oscillations. Finally, we employ a variant of the model to capture hippocampal activity during spatial navigation. Together, results highlight a novel role for neural oscillations in regulating temporal processing within recurrent networks of the brain.

## 2. Methods

### A. Integrate-and-fire networks

#### A.1. Driven networks

Our network consists of leaky integrate-and-fire neurons, where *N*_*rnn*_ = 1, 000 for the SRNN projecting to the read-out units and *N*_*osc*_ = 500 for each oscillatory network, by default. 80% of these neurons are selected to be excitatory while the remaining 20% are inhibitory. The membrane potential of all neurons is given by

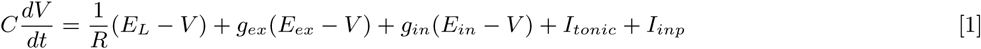

where *C* and *R* are the membrane capacitance and resistance, *E*_*L*_ is the leak reversal potential, *g*_*ex*_ and *g*_*in*_ are the time-dependent excitatory and inhibitory conductances, *E*_*ex*_ and *E*_*in*_ are the excitatory and inhibitory reversal potentials, *I*_*tonic*_ is a constant current applied to all neurons and *I*_*inp*_ is a time-varying input described below. Parameters were sampled from Gaussian distributions as described in Table 1. The excitatory and inhibitory conductances, *g*_*ex*_ and *g*_*in*_ respectively, obey the following equations:

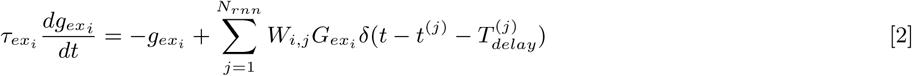

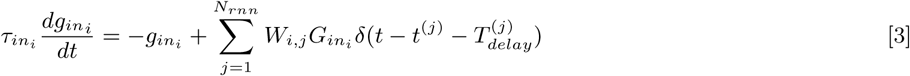

where *τ*_*ex*_ and *τ*_*in*_ are the time constants of the excitatory and inhibitory conductances, and *G*_*in*_ and *G*_*ex*_ are the change in conductance from incoming spikes to excitatory and inhibitory synapses. *V*_*θ*_ is the spiking threshold, *t*^(*j*)^ denotes the time since the last spike of the pre-synaptic neuron *j*, after which *V* is set to *V*_*reset*_ for a duration equal to *τ*_*ref*_. *T*_*delay*_ is the propagation delay of the action potential. The SRNN connectivity is defined as a *N*_*rnn*_ × *N*_*rnn*_ sparse and static connectivity matrix *W* with a density *p*_*rnn*_ (probability of having a non-zero pairwise connection). The non-zero connections are drawn from a half-normal distribution *f* (0, *σ*_*rnn*_), where 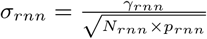. All ODEs are solved using a forward Euler method with time-step Δ*t* = 0.05 ms.

**Table 1.** SRNN parameters.

|  | Main SRNN | Input (oscillatory networks) |
| --- | --- | --- |
| N | 1,000 | 500 |
| R | 100 $M\Omega$ | 100 $M\Omega$ |
| C | 200 $pF$ | 200 $pF$ |
| $E_L$ | $\mu = -60 \text{ mV}, \sigma = 1.2 \text{ mV}$ | $\mu = -60 \text{ mV}, \sigma = 0.6 \text{ mV}$ |
| $V_\theta$ | $\mu = -50 \text{ mV}, \sigma = 0.5 \text{ mV}$ | $\mu = -50 \text{ mV}, \sigma = 0.5 \text{ mV}$ |
| $V_{reset}$ | $\mu = -60 \text{ mV}, \sigma = 1.2 \text{ mV}$ | $\mu = -60 \text{ mV}, \sigma = 0.6 \text{ mV}$ |
| $I_{tonic}$ | 90 pA | 90 pA |
| $T_{delay}$ | $\mu = 1 \text{ ms}, \sigma = 0.02 \text{ ms}$ | $\mu = 1 \text{ ms}, \sigma = 0.01 \text{ ms}$ |
| $\tau_{ref}$ | $\mu = 2 \text{ ms}, \sigma = 0.04 \text{ ms}$ | $\mu = 2 \text{ ms}, \sigma = 0.02 \text{ ms}$ |
| $G_{ex}$ | $\mu = 20 \text{ pS}, \sigma = 0.4 \text{ pS}$ | $\mu = 30 \text{ pS}, \sigma = 0.03 \text{ pS}$ |
| $G_{in}$ | $\mu = 160 \text{ pS}, \sigma = 3.2 \text{ pS}$ | $\mu = 140 \text{ pS}, \sigma = 1.4 \text{ pS}$ |
| $\tau_{ex}$ | $\mu = 20 \text{ ms}, \sigma = 0.4 \text{ ms}$ | $\mu = 20 \text{ ms}, \sigma = 0.2 \text{ ms}$ |
| $\tau_{in}$ | $\mu = 20 \text{ ms}, \sigma = 0.4 \text{ ms}$ | $\mu = 80 \text{ ms}, \sigma = 0.8 \text{ ms}$ |
| $E_{ex}$ | 0 mV | 0 mV |
| $E_{in}$ | -80 mV | -80 mV |
| $p_{rnn}$ | 0.1 | 1 |
| $p_{inp}$ | 0.3 | 1 |
| $\gamma_{rnn}$ | 1 | 1 |

Each SRNN neuron is connected to each of the *N*_*inp*_ input units with probability *p*_*inp*_. The external inputs (*I*_*inp*_) (with index *inp* dropped to alleviate notation) follow:

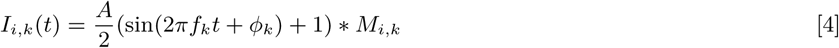

where *i* = 1, …, *N*_*rnn*_ stands for the identity of the post-synaptic unit and *k* = 1, …, *N*_*inp*_, where *N*_*inp*_ represents the total number of inputs. *f* is the frequency of the sine wave and the initial phase *ϕ* is drawn from a uniform distribution *U* (−*π, π*) (fixed for each realization of a task). The sine wave is then transformed by adding 1 and dividing by 2 to limit its range to [0,A]. The input is rescaled by the connectivity weight *M*_*i,k*_ from input unit *k* into SRNN unit *i*. The full input-to-SRNN connectivity matrix *M* is a *N*_*rnn*_ × *N*_*inp*_ sparse and static matrix, with a density of *p*_*inp*_. The non-zero connections of *M* are drawn from a normal distribution 𝒩(0, 1) and *A* is the amplitude of the input (30 pA by default).

All of the SRNN’s excitatory neurons project to the readout units. Their spiking activity *r* is filtered by a double exponential:

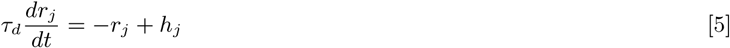

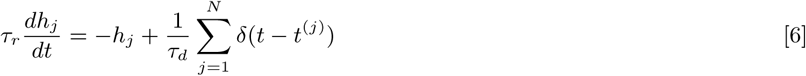

where *τ*_*r*_ = 6 ms is the synaptic rise time and *τ*_*d*_ = 60 ms is the synaptic decay time. *W*_*out*_ is initialized as a 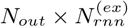 null matrix that is modified according to the learning rule described below (see training procedure). *N*_*out*_ is the number of readout units and 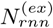 is the number of excitatory neurons in the SRNN.

### B. Target functions

Unless otherwise stated, the target functions employed to train the model were generated from white noise with a normal distribution *N* (0, 30), then low-pass filtered with a cut-off at 6 Hz. To assess network performance, we computed the Pearson correlation between the output of the network and the target function.

### C. Learning algorithm for the readout unit

We used the recursive least square algorithm (20) to train the readout units to produce the target functions. The *W*_*out*_ weight matrix was updated based on the following equations:

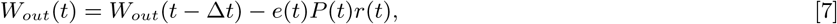

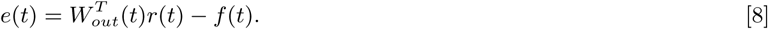

Where the error *e*(*t*) was determined by the difference between the value of the readout unit obtained with the multiplication of the reservoir’s activity with the weights *W*_*out*_, and the target function’s value *f* at time *t*. Each weight update was separated by a time interval Δ*t* of 2.5 ms for all simulations. *P* is a running estimate of the inverse of the correlation matrix of the network rates *r* (see eq. Eq. (5)), modified according to eq. Eq. (9) and initialized with eq. Eq. (10).

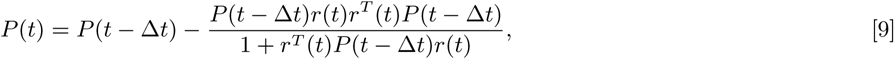

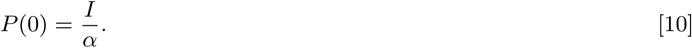

where *I* is the identity matrix and *α* is a learning rate constant.

### D. Oscillatory networks

#### D.1. External drive

Oscillatory networks obey the same equations as the SRNNs. However, the *I*_*inp*_ term in eq.Eq. (1) is replaced by the following step functions:

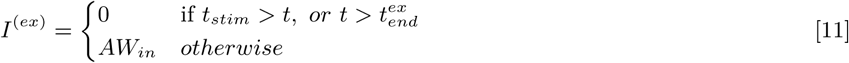

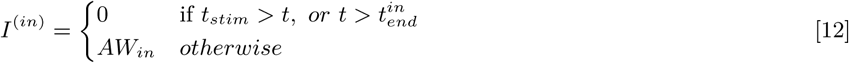

where *t*_*stim*_ = 500 ms, 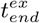 denotes the end of the excitatory pulse and 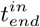 (different across oscillatory networks) is the end of the inhibitory pulse, where 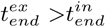. *W*_*in*_ is a dense matrix with *N*_*osc*_ × *N*_*inp*_ elements representing the connections from the tonic inputs to the oscillatory networks, where *N*_*inp*_ is the number of oscillatory networks projecting to the SRNN and *A* = 20 pA.

The oscillatory networks each project to the SRNN with a *N*_*rnn*_ × (*N*_*osc*_ × *N*_*inp*_) sparse and static connectivity matrix *M*, where each oscillatory network projects to the SRNN with a density of *p*_*inp*_ = 0.5. The non-zero connections are drawn from a normal distribution 𝒩 (0, *σ*_*inp*_), where:

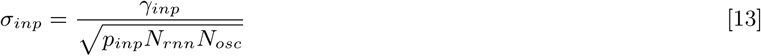

and *γ*_*inp*_ = 10 (a.u.) by default.

Where stated in the Results, the SRNN projects back to the oscillatory networks with a (*N*_*osc*_ × *N*_*inp*_) × *N*_*rnn*_ sparse and static connectivity matrix *M*′, where each SRNN unit projects to the oscillatory networks with a density of *p*_*fb*_ = 0.5. The non-zero connections are drawn from a normal distribution 𝒩 (0, *σ*_*fb*_), where:

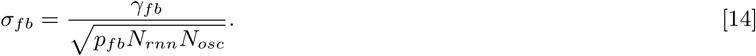

#### D.2. Selection of stable networks

As we were interested in regimes where the networks would produce reliable and repeatable oscillations to be used as an input to our model, we considered networks with an inter-trial correlation coefficient (10 trials) of their mean firing-rate greater than 0.95 as stable. A wide range of parameter combinations lead to reliable oscillations, but different random initialisations of networks with the same parameters can lead to drastically different behavior, both in activity type (ansynchronous and synchronous) and inter-trial reliability (Fig. S3).

### E. Jitter accumulation in input phase

While a perfect sinusoidal input such as the one in eq. Eq. (21) allows for well-controlled simulations, it is unrealistic from a biological standpoint. To address this issue, we added jitter to the input phase of each input unit. This was achieved by converting the static input phase injected into unit k *ϕ*_*k*_ to a random walk *ϕ*_*k*_(*t*). First, we discretize time into non-overlapping bins of length Δ*t*, such that *t*_*n*_ = Δ*t* ∗ *n*. From there, we iteratively define *ϕ*(*t*) (index *k* is dropped to alleviate the notation) as:

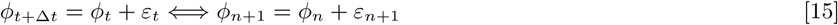

with

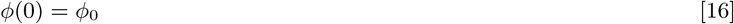

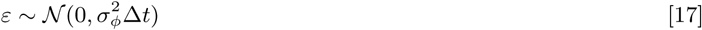

from initial value *ϕ*_0_, sampled from a uniform distribution as specified previously. More intuitively, *ϕ*(*t*) can be constructed as:

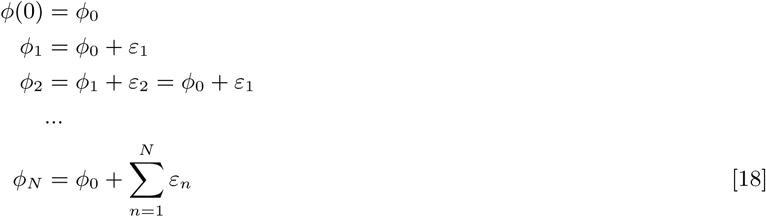

On average, the resulting deviation from the deterministic signal, *i.e. E* [*ϕ*_*N*_ − *ϕ*_0_], is null. On the other hand, one can calculate its variance:

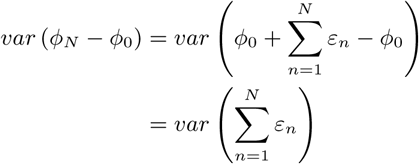

Since all *ε*_*n*_ are i.i.d:

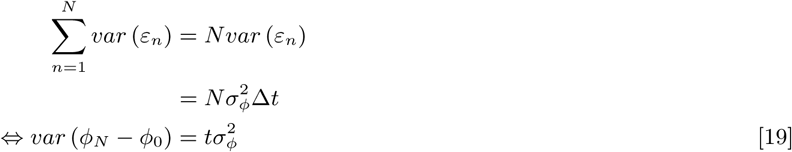

For ease of comparison, we can express the equivalent standard deviation in degrees (see Fig. S7):

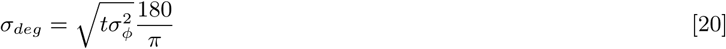

### F. Model for place cells sequence formation

#### F.1. Network architecture and parameters

We employed a balanced recurrent network similar to the ones used for all other simulations, with a few key differences. The input consisted of *N*_*inp*_ = 20 oscillators with periods ranging from 7.5 to 8.5 Hz that densely projected to the SRNN (*p* = 1) and follow:

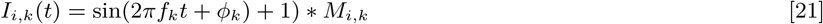

*C* was set to 100 pF for all neurons and *γ*_*rnn*_ was set to 0.5. We removed the readout unit and connections, and we selected 10 random excitatory cells (*N*_*place*_) as place cells. Those cells had parameters identical to the other SRNN excitatory units, except:

1. We set the resting potential of those cells to the mean of *E*_*L*_, to avoid higher values that could lead to high spontaneous activity (that in turn can lead to spurious learning).
2. A 600 ms sine wave at 10 Hz with an amplitude of 60 pA was injected in each of the place cells at a given time representing the animal going through its place field.
3. The connections between the input oscillators and the place cells were modified following eq.Eq. (22).

We modelled the environmental input (10 Hz depolarisation of CA1 place cells) based on a representation of the animal’s location that was fully dependent on time. In order to explain phase precession, our model relied on an environmental input of a slightly higher frequency than the background theta oscillation, as suggested in (21).

The learning rule seeks to optimize the connections between the oscillating inputs and the place cells in order to make them fire whenever the right phase configuration is reached (18, 19).

We used a band-pass filter between 4-12 Hz to isolate the theta rhythm in the SRNN. We then used a Hilbert transform to obtain the instantaneous phase of the resulting signal.

#### F.2. Learning algorithm

We developed a correlation-based learning rule (22) inspired by the results obtained by (23):

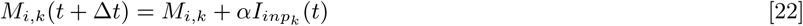

where *i* = 1, …, *N*_*place*_ and *α* = 0.25. With this rule, the weight update is only applied when a burst occurs in the place cells. A burst is defined as any spike triplets that occur within 50 ms. In experiments, these post-synaptic bursts were associated with *Ca*^2+^ plateaus in place cells (23) that lead to a large potentiation of synaptic strength with as few as 5 pairings. The connections of *M* were initialized from a half-normal distribution *f* (0, *σ*_*inp*_), where *σ*_*inp*_ = 0.1 and the signal amplitude *A* was set to 1. *M* was bound between 0 and 5\**σ*_*inp*_ during training.

### G. Audio processing for speech learning

We used the numpy/python audio tools from (24), adapted by (25), to process the audio WAVE file. We used the built-in functions to convert the audio file to a mel-scaled spectrogram and to invert it back to a waveform.

## 3. Results

### A. A cortical network driven by oscillations

We began with a basic implementation of our model where artificial oscillations served as input to a SRNN (Fig. 1a) – in a later section, we will describe a more realistic version where recurrent networks generate these oscillations intrinsically. In this simplified model, two input nodes, but potentially more (Fig. S1), generate sinusoidal functions of different frequencies. These input nodes project onto a SRNN that is a conductance-based leaky integrate-and-fire (LIF) model (26) with balanced excitation and inhibition (7). Every cell in the network is either strictly excitatory or inhibitory, thus respecting Dale’s principle.

**Fig. 1.**
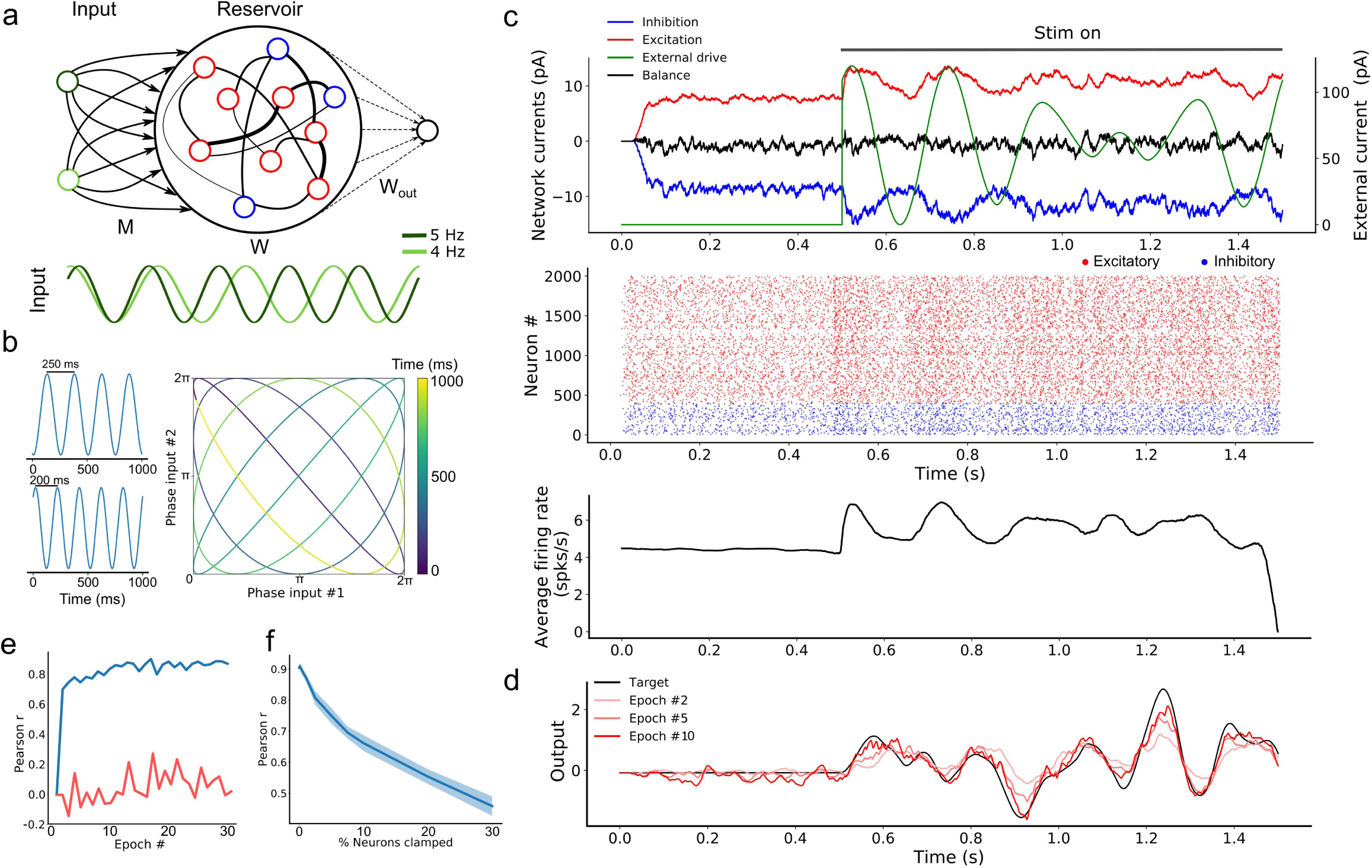
Oscillation driven SRNN to learn complex temporal tasks. **a** Schematic of the model’s architecture. This implementation has two input units that inject sine-waves in a subset of the SRNN’s neurons. *M* denotes the connections from the input units to the SRNN neurons. *W* denotes the recurrent connectivity matrix of the SRNN. *W*_*out*_ denotes the trainable connections from the excitatory SRNN neurons to the readout unit. **b** Example showing how two sine waves of different frequencies can be combined to generate a two-dimensional function with a period longer than either sine wave. **c** Sample average current per neuron, raster and average instantaneous firing rate over one trial, respectively. The external drive is only delivered to a subset of SRNN units (*p*_*inp*_ = 0.3) for each input unit. **d** Output of the readout unit during testing trials (learning rule turned off) interleaved during training. The nearly constant output without the input is due to random activity in the SRNN **e** Pearson correlation between the output of the network and the target function as a function of the number of training epochs when driven with (blue) and without (red) multi-periodic input. **f** Pearson correlation between the output and the target function as neurons of the SRNN are clamped.

The combination of *N*_*inp*_ input oscillators will generate a sequence of unique *N*_*inp*_-dimensional vectors where the sequence lasts as long as the least common multiple of the inputs’ individual periods (14). For instance, two sine waves with periods of 200 ms and 250 ms would create a multi-periodic input with a period lasting 1,000 ms. This effect can be viewed as a two-dimensional state-space where each axis is an individual sine wave (Fig. 1b). Using a phase reset of the oscillations on every trial can then evoke a repeatable pattern of activity in the downstream population of neurons. Thus, multiplexed oscillations provide the network with inputs whose timescale largely exceeds that of individual units.

When a SRNN (*N*_*res*_ = 2,000) was injected with oscillations, excitatory and inhibitory populations modulated their activity over time, while the average input currents to individual neurons remained balanced (Fig. 1c, top panel). To illustrate the benefits of oscillatory inputs on a SRNN, we designed a simple task where a network was trained to reproduce a target function consisting of a time-varying signal generated from low-pass filtered noise (Fig. 1c, bottom panel). Simulations were split into a training and a testing phase. During the training phase, the network was training when receiving a combination of two oscillatory inputs at 4 and 5 Hz. Synaptic weights from the SRNN to the read-out were adjusted using the recursive least-squares learning algorithm (20) adapted to spiking units (13). During the testing phase, synaptic weights were frozen and the network’s performance was assessed by computing the Pearson correlation between the target function and the network’s output. This correlation increased to 0.9 within the first 10 training epochs and remained stable thereafter (Fig. 1d). By comparison, the output of a similar network with no oscillatory inputs remained uncorrelated to the target function. Thus, oscillatory inputs create rich and reliable dynamics in the SRNN that enabled the read-out to produce a target function that evolved over time in a precise manner.

Next, we investigated the resilience of the network to structural perturbations where a number of individual neurons from the SRNN were “clamped” (i.e., held at resting potential) after training (14). We trained a network for 10 epochs, then froze the weights and tested its performance on producing the target output. We then gradually clamped an increasing proportion of neurons from the SRNN. The network’s performance decreased gradually as the percentage of clamped units increased (Fig. 1e). Remarkably, the network produced an output that correlated strongly with the target function (correlation of 0.7) even when 10% of neurons were clamped. Further exploration of the model shows a wide range of parameters that yield high performances (Fig. S1). Oscillatory inputs thus enabled SRNN to produce precise and repeatable patterns of activity under a wide range of modeling conditions. Next, we improved upon this simple model by developing a more biologically-inspired network that generated oscillations intrinsically.

### B. Endogenously generated oscillatory activity

While our results thus far have shown the benefits of input oscillations when training a SRNN model, we did not consider their neural origins. To address this issue, we developed a model that replaces this artificial input with activity generated by an “oscillator” spiking network acting as a central pattern generator (27).

To do so, we took advantage of computational results showing that sparsely connected networks can transition from an asynchronous to a periodic synchronous regime in response to a step current (15, 28, 29), thus capturing *in vivo* activity (30) (see Methods) (Fig. 2a). The periodicity of the synchronous events could therefore potentially be used to biologically capture the effects of artificially generated sine waves.

**Fig. 2.**
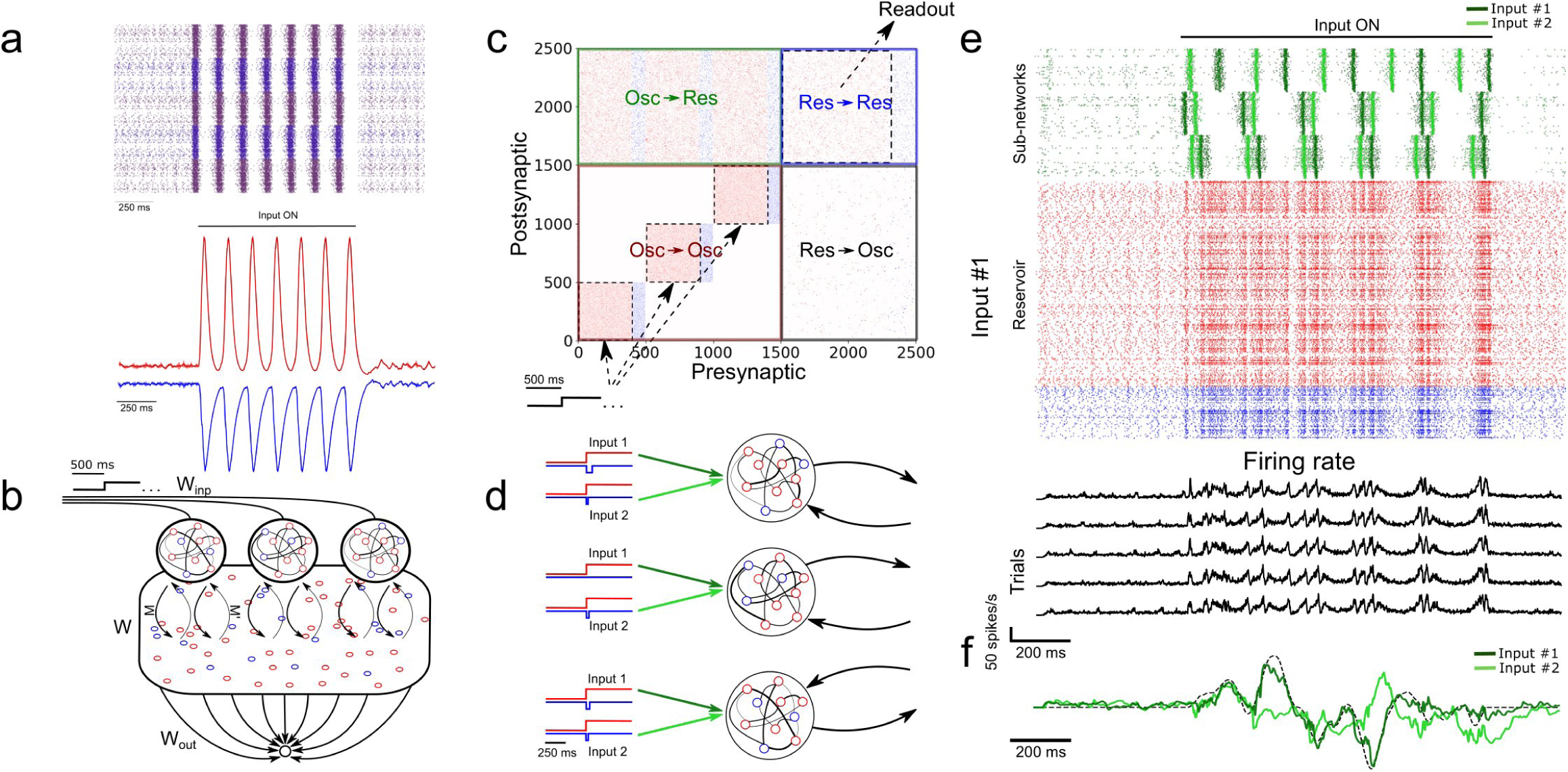
SRNN driven with intrinsically generated oscillations. **a** Top: Sample rasters from one network on five different trials. Bottom: Average and standard error of the excitatory (red) and inhibitory (blue) conductance of a stable network on 5 different trials (see Methods). The network is asynchronous when the external drive is off, and becomes periodic when turned on. **b** Architecture of the augmented model. As in 1a, *W* and *W*_*out*_ denote the recurrent and readout connections, respectively. *M* is the connection matrix from the input to the SRNN, except that the input units are now replaced by networks of neurons. *M′* denotes the feedback connections from the SRNN to the oscillatory network. *W*_*inp*_ denotes the connections providing the tonic depolarization to the oscillatory networks. **c** Connectivity matrix of the model. The external drive is provided solely to the oscillatory networks that project to the SRNN that in turn projects to the readout unit. **d** External inputs provided to the oscillatory networks with varied inhibitory transients associated with each excitatory input. **e** Top: Sample activity of the oscillatory networks (green) on two separate trials with different inputs, and the SRNN for one trial (input #1). Bottom: PSTH of the network’s neurons activity on five different trials with input #1. **f** Post-training output of the network, where input #1 was paired with the target, but input #2 was not.

In simulations, we found that this transition was robust to both synaptic noise and neuronal clamping (Fig. S2). Further, the frequency of synchronized events could be modulated by adjusting the strength of the step current injected in the network, with stronger external inputs leading to a higher frequency of events (Fig. S2). Thus, oscillator networks provide a natural neural substrate for input oscillations into a recurrent network.

From there, we formed a model where three oscillator networks fed their activity to a SRNN (Fig. 2b). These oscillator networks had the same internal parameters except for the inhibitory decay time constants of their recurrent synapses (*τ*_*in*_ = 70, 100 and 130 ms for each network) thus yielding different oscillatory frequencies (Figs. S3,S4). In order to transition from an asynchronous to a synchronous state, the excitatory neurons of the oscillator networks received a step current.

The full connectivity matrix of this large model is depicted in Fig. 2c. As shown, the oscillator networks send sparse projections (with a probability of 0.3 between pairs of units) to the SRNN units. Only the excitatory neurons of the SRNN project to the readout units. Only weak feedback from the SRNN to the oscillators was included – in supplementary simulations, we found that strong feedback projections desynchronized the oscillator networks (Figs. S2,S5).

A sample of the full model’s activity is shown in Fig. 2e. Both the oscillator and the SRNN showed asynchronous activity until a step current was injected into the excitatory units of the oscillator networks. In response to this step current, both oscillator and SRNN transitioned to a synchronous regime. The model reverted back to an asynchronous regime once the step current was turned off.

To illustrate the behavior of this model, we devised a “cued” task similar to the one described above, where the goal was to reproduce a random time-varying signal. When learning this signal, however, the oscillator networks received a cue (“Input 1”) consisting of a combination of excitatory step current and transient inhibitory input (Fig. 2d) that alters the relative phase of the input oscillators, but not their frequency. Following 20 epochs of training, we switched to a testing phase and showed that the model closely matched the target signal (Fig. 2f). Crucially, this behavior of the model was specific to the cue provided during training: when a different, novel cue (shaped by inhibitory transients) was presented to the network (“Input 2”), a different output was produced (Fig. 2f).

In sum, the model was able to learn a complex time-varying signal by harnessing internally-generated oscillations that controlled the ongoing activity of a SRNN. In the following section, we aimed to further explore the computational capacity of the model by training a SRNN on multiple tasks in parallel.

### C. An artificial network that learns to multitask

To explore the ability of the model to learn two tasks concurrently, we reverted to our initial model with artificial oscillations, allowing for a more principled control of the input injected to the SRNN. The oscillatory input consisted of three sine waves of different frequencies (Fig. 3a) (more inputs lead to richer dynamics, Fig. S1).

**Fig. 3.**
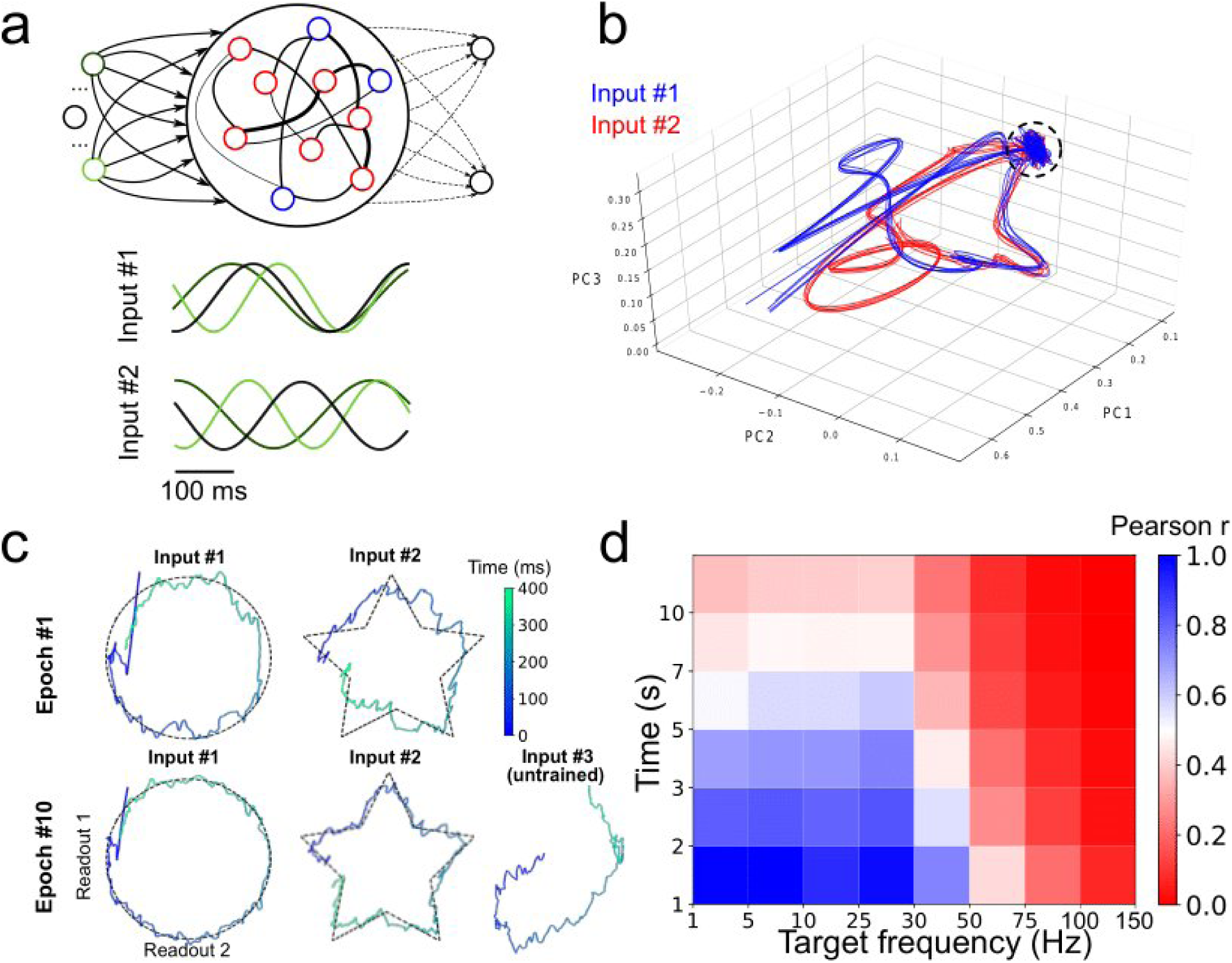
Parallel training of multiple tasks. **a** Schematic of the model’s architecture with an additional readout unit. An example of possible phase shifted inputs is shown at the bottom. **b** Trajectory of the SRNN on the first three components of a PCA. Without an external input, the network is spontaneously active and constrained in a subspace of the state-space (depicted by the circle). Upon injection of the input, the SRNN’s activity is kicked into the trajectory related to the input. **c** Output of the two readout units after the first and tenth training epochs. **d** Heat map showing the performance of the model on tasks with different lengths and frequencies.

We trained this model on two different motor control tasks that required the network to combine the output of two readout units in order to draw either a circle or a star in two dimensions. Here, each of the outputs corresponded to x- and y-coordinates, respectively. The phase of the oscillations (“Input 1” vs. “Input 2”) were individually paired with only one of the two tasks in alternation (Fig. 3a).

We employed a principal component analysis (PCA) to visualize the activity of the SRNN before and during training (Fig. 3b). Before injecting the oscillatory inputs, the network generated spontaneous activity that occupied a limited portion of the state space (Fig. 3b, circle). During training, the oscillatory inputs were turned on, resulting in different trajectories depending on the relative phase of the oscillations. The network thus displayed a distinct pattern of activity for each of the two tasks.

Viewed in two dimensions, the outputs of the network rapidly converged to a circle and a star that corresponded to each of the two target shapes when given each respective input separately (Fig. 3c). These shapes were specific to the particular phase of the oscillatory input – in a condition where we presented a randomly-chosen phase configuration to the network, the output did not match either of the trained patterns (Fig. 3c).

Finally, we tested the ability of the model to learn a number of target signals varying in duration and frequency. We generated a number of target functions consisting of filtered noise (as described previously), and varied their duration as well as the cut-off frequencies of band-pass filtering. The network performed optimally for tasks with relatively low frequency (<30 Hz) and shorter duration (<5 seconds), and had a decent performance for even longer targets (e.g, correlation of *r*=0.5 for a time of 7 seconds and a frequency below 30 Hz). (Fig. 3d).

In sum, the model was able to learn multiple tasks in parallel based on the phase configuration of the oscillatory inputs to the SRNN units. The range of target signals that could be learned was dependent upon their duration and frequency. The next section will investigate another aptitude of the network, where a target signal can be rescaled in time without further training.

### D. Temporal rescaling of neuronal activity

A key aspect of many behavioral tasks based on temporal sequences is that once learned they can be performed faster or slower without additional training. For example, when a new word is learned, it can be spoken faster or slower without having to learn the different speeds separately.

We propose a straightforward mechanism to rescale a learned temporal sequence in the model. Because the activity of the model strongly depends upon the structure of its oscillatory inputs, we conjectured that the model may generate a slower or faster output by multiplying the period of the oscillatory inputs by a common factor. Biologically, such a factor might arise from afferent neural structures that modulate oscillatory activity (15). Due to the highly non-linear properties of the network, it is not trivial that rescaling the inputs would expand or compress its activity in a way that preserves key features of the output (31).

To test the above mechanism, we trained a SRNN receiving sine wave inputs to produce a temporal sequence of low-pass filtered random activity. After the pair of input-target was trained for 10 epochs, we tested the network by injecting it with sine waves that were either compressed or expanded by a fixed factor relative to the original inputs (Fig. 4a). To evaluate the network’s ability to faithfully replay the learned sequence, we computed the Pearson correlation between the output of the network (Fig. 4b) and a compressed or expanded version of the target signal.

**Fig. 4.**
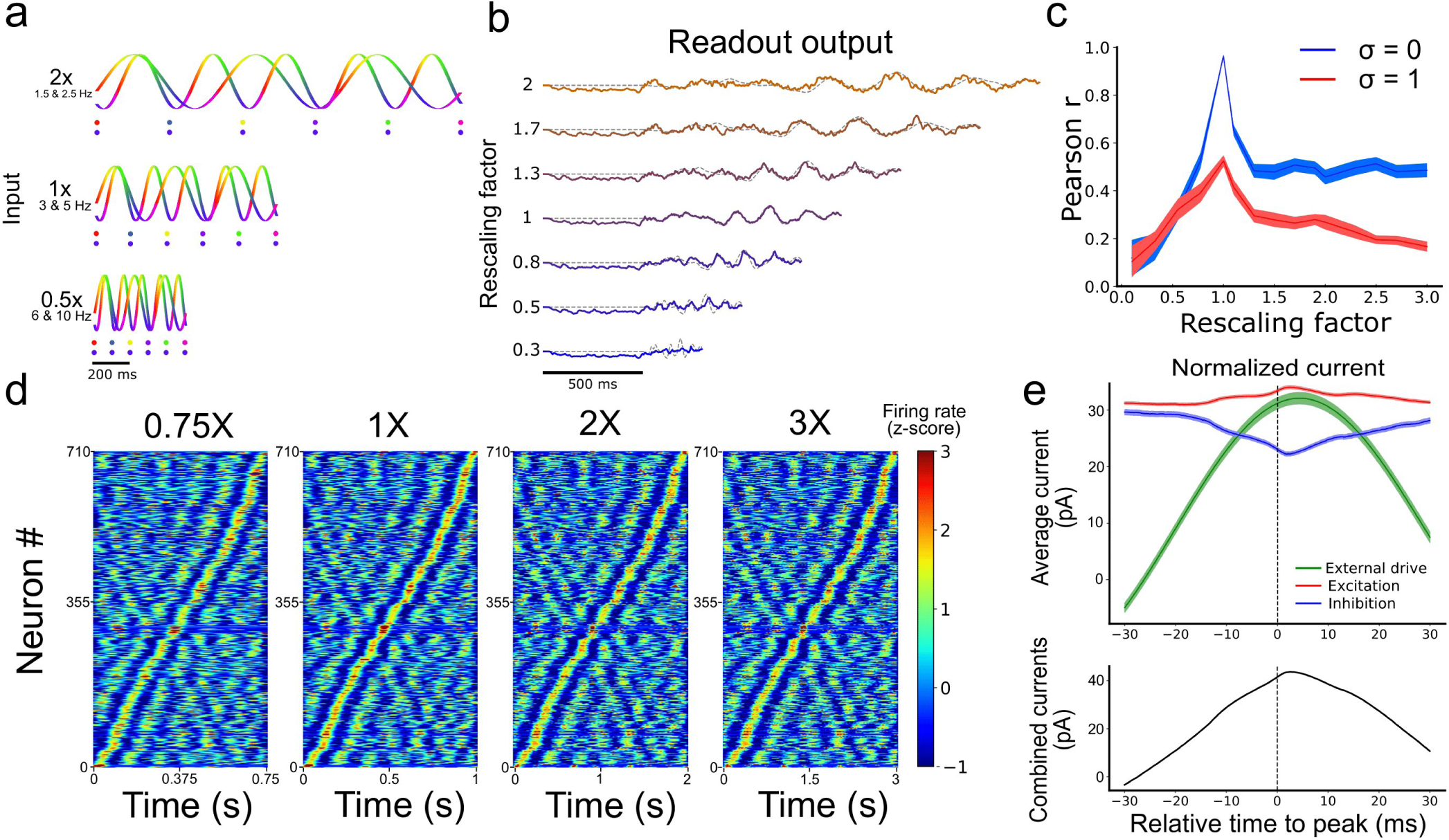
Temporal rescaling of the network’s activity. **a** Activity of two input units with different rescaling factors. Each phase is represented in a different color. The two dots shown for the different velocities are spaced at a constant proportion of the whole duration, and show that the phase alignment of the two oscillators is preserved across different rescaling factors. **b** Output of the network for different rescaling factors. The target is represented by the grey dashed line. **c** Pearson correlation between the output and the target function for different rescaling factors and different input noise variance (*σ*). **d** Spiking activity of the RNN at 0.75X, 1X, 2X, and 3X sorted based on the peak of the activity at the original scaling (1X). **e** Average excitatory (red), inhibitory (blue), and external input (green) received by each cell aligned on their peak firing rate (at t = 0 ms). The combined inputs (black, external + excitation – inhibition) is represented at the bottom.

Performance degraded gradually with inputs that were expanded or contracted in time relative to the target signal (i.e, as the rescaling factor moved further away from 1) (Fig. 4c). Further, performance degraded more slowly beyond a rescaling factor of 1.5, particularly when input noise was absent, suggesting some capacity of the network to expand the target signal in time (Fig. S6). This result offered a qualitative match to experimental findings (32) and the performance of the network was tolerant to small phase deviations resulting from the addition of random jitter in the phase of the input oscillations (Fig. S7).

In sum, rescaling the speed of the input oscillations by a common factor lead to a corresponding rescaling of the learned task, with compressed neural activity resulting in more error than expanded activity. The next section examines some of the underlying features of activity in a SRNN driven by oscillatory inputs.

### E. Temporal selectivity of artificial neurons

A hallmark of temporal processing in brain circuits is that some subpopulations of neurons increase their firing rate at specific times during the execution of a timed task (33–35). To see whether this feature was present in the model, we injected similar oscillatory inputs as above (10 input units between 5 and 10 Hz for 1 second) for 30 trials. To match experimental analyses, we convoluted the firing rate of each neuron from the SRNN with a Gaussian kernel (s.d. = 20 ms), averaged their activity over all trials, and converted the resulting values to a z-score. To facilitate visualization, we then sorted these z-scores by the timing of their peak activity. We retained only the neurons that were active during the simulations (71%). Results showed a clear temporal selectivity whereby individual neurons increased their firing rate at a preferred time relative to the onset of each trial (Fig. 4d).

These “selectivity peaks” in neural activity were maintained in the same order when we expanded or contracted the input oscillations by a fixed factor (Fig. 4a), and sorted neurons based on the original input oscillations (Fig. 4d), thus capturing recent experimental results (34). To shed light on the ability of simulated neurons to exhibit temporal selectivity, we examined the timing of excitatory and inhibitory currents averaged across neurons of the SRNN. We then aligned these currents to the timing of selectivity peaks and found elevated activity around the time of trial-averaged peaks (Fig. 4e). Therefore, both the input E/I currents and the external inputs drive the activity of the neurons near their peak response, showing that both intrinsic and external sources drives the temporal selectivity of individual neurons.

In sum, neurons from the SRNN show sequential patterns of activity by leveraging a combination of external drive and recurrent connections within the network. Next, we examined the ability of the model to learn a naturalistic task of speech production.

### F. Learning Natural Speech, Fast and Slow

In a series of simulations, we turned to a biologically and behaviorally relevant task of natural speech learning. This task is of particular relevance to temporal sequence learning given the precise yet flexible nature of speech production: once we learn to pronounce a word, it is straightforward to alter the speed at which this word is spoken without the need for further training. We thus designed a task where an artificial neural network must learn to utter spoken words in the English language and pronounce them slower or faster given the appropriate input, without retraining.

To train a network on this task, we began by extracting the waveform from an audio recording of the word “reservoir” and converting this waveform to a spectrogram (Fig. 5a). We then employed a compression algorithm to bin the full range of frequencies into 64 channels spanning a range from 300 Hz to 8 kHz (see Methods). Each of these channels were mapped onto an individual readout unit of the model. Synaptic weights of the SRNN to the readout were trained to reproduce the amplitude of the 64 channels over time. The output spectrogram obtained from the readout units was converted to an audio waveform and compared to the target waveform (see Movie S1).

**Fig. 5.**
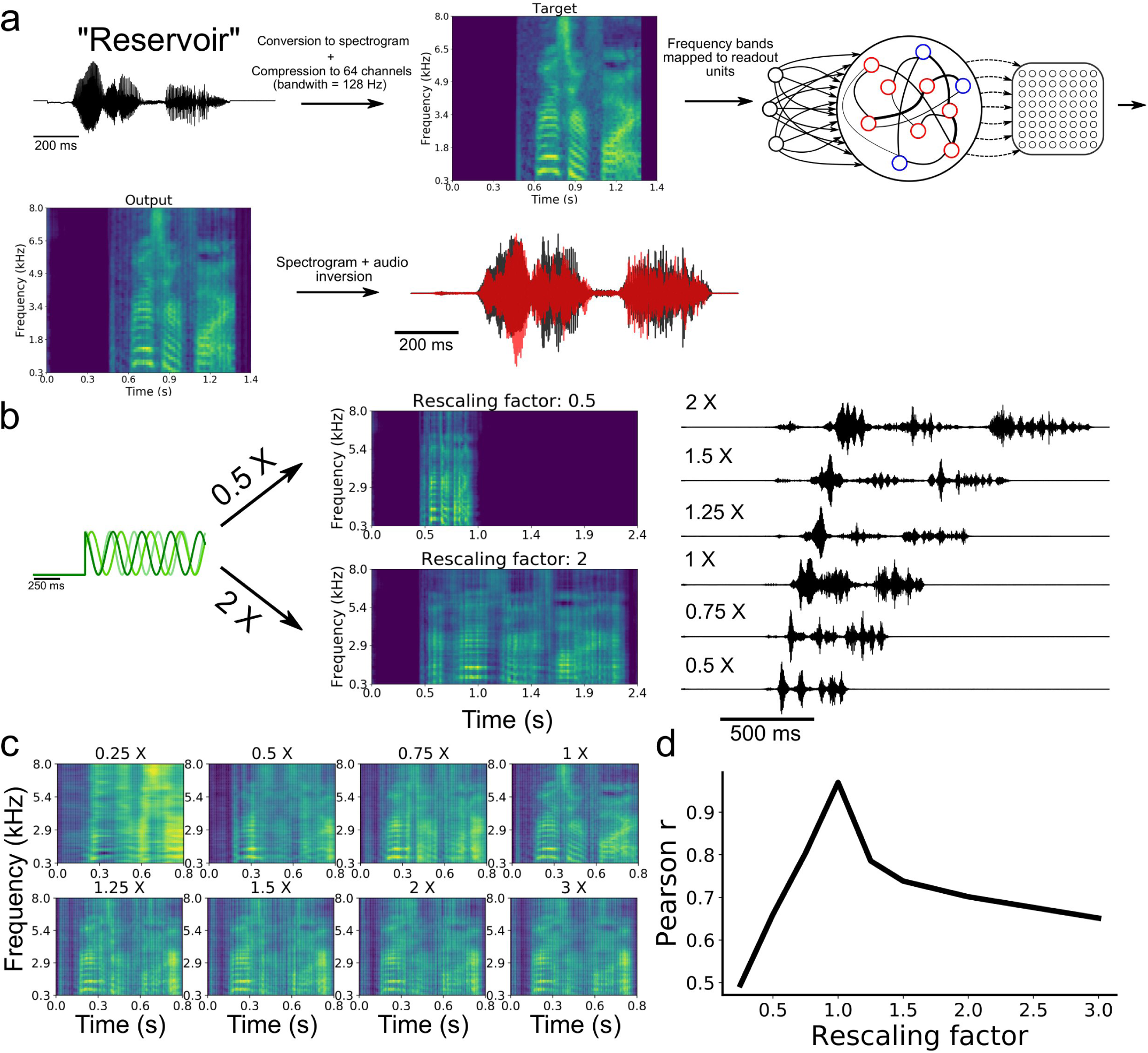
Speech learning and production with temporal rescaling. **a** Workflow of the transformation and learning of the target audio sequence. **b** The input frequencies are either sped-up or slowed-down in order to induce temporal rescaling of the speed of execution of the task. **c** The rescaled output spectrograms are scaled back to the original speed in order to compare them to the target spectrogram. **d** Average correlation between the output and target of all channels of the spectrograms for the different rescaling factors.

Following training, the network was able to produce a waveform that closely matched the target word (Fig. 5a). To examine the ability of the network to utter the same word faster or slower, we employed the rescaling approach described earlier, where we multiplied the input oscillations by a constant factor (Fig. 4a).

Our model was able to produce both faster and slower speech than what it had learned (Fig. 5b). Scaling the outputs back to the original speed showed that the features of the spectrogram were well replicated (Fig. 5c). The correlation between the rescaled outputs and target signal decreased as a function of the rescaling factor (Fig. 5d), in a manner similar to the above results on synthetic signals (Fig. 4c).

Results thus suggest that multiplexing oscillatory inputs enabled a SRNN to acquire and rescale temporal sequences obtained from natural speech. In the final section below, we employed our model to capture hippocampal activity during a well-studied task of spatial navigation.

### G. Temporal sequence learning during spatial navigation

Thus far, we have modelled tasks where units downstream of the SRNN are learning continuous signals in time. In this section, we turned to a task of spatial navigation that required the model to learn a discrete sequence of neural activity.

A wealth of experiments shows that subpopulations of neurons become selectively active for specific task-related time intervals. A prime example is seen in hippocampal theta sequences (36) that are observed during spatial navigation in rodents, where individual place cells (37) increase their firing rate at a given location in space (place fields). During spatial navigation, the hippocampus shows oscillatory activity in the theta range (4-12 Hz), likely originating from both the medial septum (38) and within hippocampus (39, 40).

In a series of simulations, we examined how neurons in a SRNN may benefit from theta oscillations to bind and replay such discrete place cells sequences. We designed a SRNN (associated with area CA1,(36)) that received inputs from multiple input oscillators (CA3, (41)) (Fig. 6a,b). We randomly selected 10 excitatory units within the SRNN and labelled them as “place cells”. To simulate the response of place cells to an environmental input indicating the spatial location of the animal (42), we depolarized these cells by an oscillating input at 10 Hz for 600 ms with a specific onset that differed across neurons in order to capture their respective place fields (assuming a fixed spatiotemporal relation of 100 ms = 5 cm on a linear track). In this way, the sequential activation of place cells from the SRNN mimicked the response of CA1 neurons to an animal walking along a linear track (Fig. 6c).

**Fig. 6.**
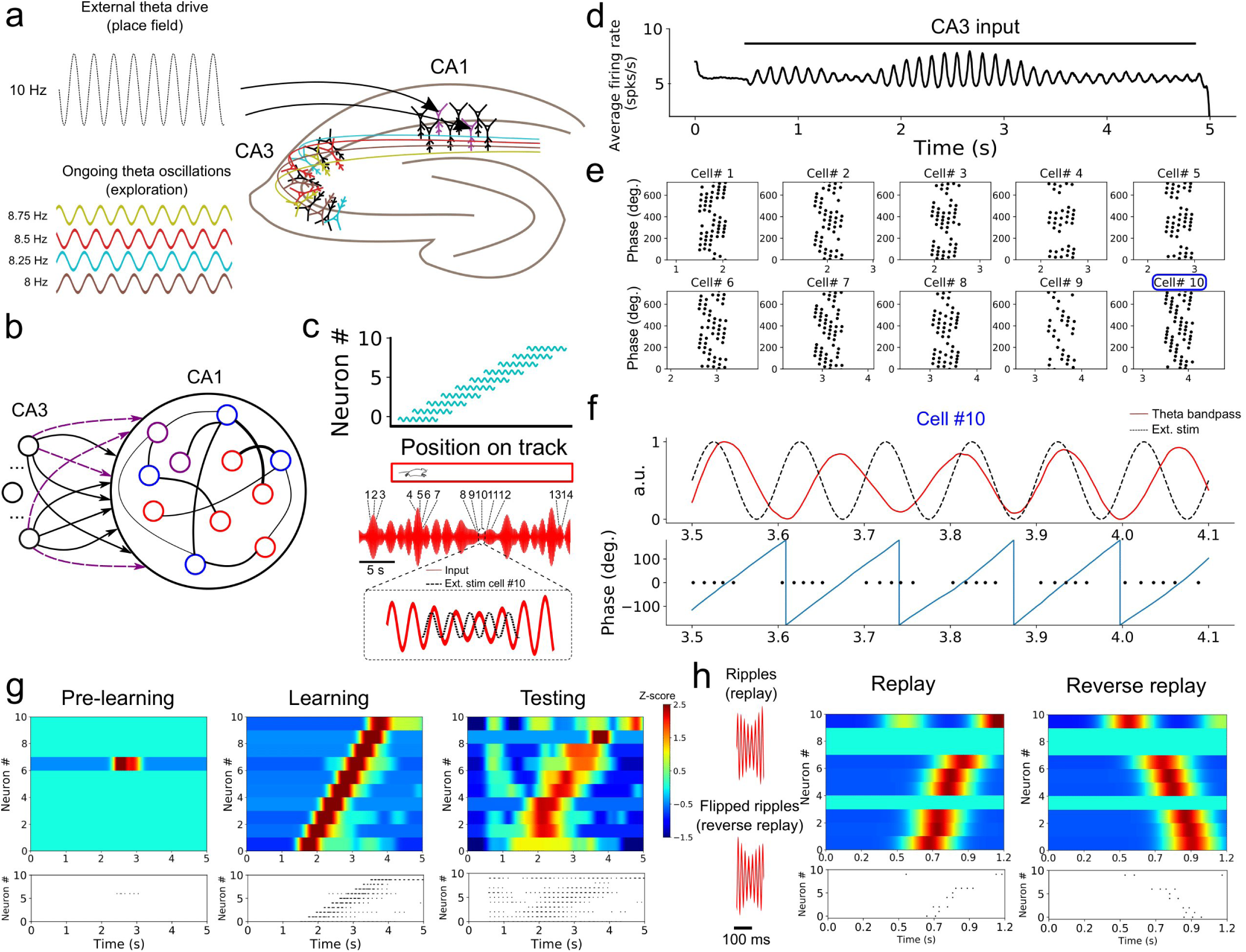
Formation of place cell sequences and replay. **a** Subpopulations of CA3 cells oscillating in the theta range and projecting to CA1. CA1 place cells are driven by a slightly faster oscillating input upon entering their place field. **b** RC implementation of the phenomenological model. The input layer is composed of oscillatory units (CA3) and CA1 is modelled by a SRNN where 10 excitatory units (purple) are randomly selected as place cells. The connections from the input to the place cells are subject to training. **c** Top: Each of the 10 SRNN place cells were driven by a depolarizing oscillating input in a temporal sequence analogous to a mouse moving along a linear track. Bottom: Resulting theta frequency of the combined oscillations (red). Each number shows a place cell activated at a given time along the ongoing theta input. The multi-periodic input from CA3 guarantees that each place cell is activated with a unique combination of the input, following the sequence in which the cell is active. **d** Multiplexing CA3 inputs generates a visible theta oscillation in the CA1 SRNN. **e** Spike times of the ten place cells in relation to the phase of the population theta activity. Each dot represents a spike at a certain time/position. Each cell shows a shift towards earlier theta phase as the animal moves along its place fields. **f** Activity of cell #3 upon entering its place field. Top: theta band-passed activity of the SRNN (red) and place field related input (dashed black). Bottom: phase of the theta oscillation (grey) and spike times. **g** Top shows a heatmap of the place cell activity and bottom shows the spike raster during each phase of training. Before training: All place cells are silent. During training: place cells are depolarized upon entering their place field. After training: a similar sequence is evoked without the external stimulation used during training. **h** Rescaling the input (factor of X0.15) leads to a high-frequency input reminiscent of ripples. A compressed version of the sequence learned in **g** is evoked, and a reversed sequence is evoked when a reverse “ripple” is injected in the SRNN.

To capture the effect of theta oscillations on CA1 activity, all neurons from the SRNN were driven by a combination of multiple oscillating inputs where the frequency of each input was drawn from a uniform distribution in the range of 7-9 Hz. Connections from the input units and the place cells within the SRNN were modified by a synaptic plasticity rule (see Methods).

As expected from the input oscillators, mean population activity of the SRNN exhibited prominent theta activity (Fig. 6d). To assess the baseline performance of the model, we ran an initial simulation with oscillatory inputs but no synaptic plasticity or place fields (i.e., no environmental inputs to the place cells). All place cells of the model remained silent (Fig. 6g). Next, we ran a training phase simulating a single lap of the virtual track lasting 5 seconds, where place cells received oscillatory inputs (CA3) as well as a depolarizing oscillation (10 Hz) whenever the cell entered its place field. During this lap, individual place cells entered their respective field only once. We assessed the performance of the model during a testing phase where both synaptic plasticity and depolarizing oscillations were turned off. During the testing phase, place cells yielded a clear sequence of activation that matched the firing pattern generated during training (Fig. 6g). Thus, place cell activity was linked to the phase of the oscillatory inputs after a single lap of exploration.

Going further, we explored two key aspects of place cell activity in the SRNN that are reported in hippocampus, namely phase precession and rapid replay. During phase precession, the phase of firing of place cells exhibits a lag that increases with every consecutive cycle of the theta oscillation (36, 43). We examined this effect in the model by extracting the instantaneous phase of firing relative to the global firing rate filtered between 4-12 Hz. The activity of individual place cells from the SRNN relative to theta activity exhibited an increasing phase lag characteristic of phase precession (Fig. 6e,f).

A second feature of hippocampal activity is the rapid replay of place cells during rest and sleep in a sequence that mirrors their order of activation during navigation (44). This replay can arise in either a forward or reverse order from the original sequence of activation (45). We compressed (factor of 0.15) the CA3 theta oscillations injected in the SRNN during training, resulting in rapid (50-55 Hz) bursts of activity (Fig. 6h). These fast oscillations mimicked the sharp-wave ripples that accompany hippocampal replay (44). In response to these ripples, place cells of the SRNN exhibited a pattern of response that conserved the order of activation observed during training (Fig. 6h). Further, a reverse replay was obtained by inverting the ripples (that is, reversing the order of the compressed sequence) presented to the SRNN (Fig. 6h).

In sum, oscillatory inputs allowed individual neurons of the model to respond selectively to external inputs in a way that captured the sequential activation and replay of hippocampal place cells during a task of spatial navigation.

## 4. Discussion

### A. Summary of results

Taken together, our results suggest that large recurrent networks can benefit from autonomously generated oscillatory inputs in order to learn a wide variety of artificial and naturalistic signals, and exhibit features of neural activity that closely resemble neurophysiological experiments.

One series of simulations trained the model to replicate simple shapes in 2D coordinates. Based upon the structure of its oscillatory inputs, the model flexibly switched between two shapes, thus showing a simple yet clear example of multitasking with a recurrent network.

When we modulated the period of input oscillations delivered to neurons of the SRNN, the model was able to produce an output that was faithful to the target signal, but sped up or slowed down by a constant factor (32, 34). Oscillations served to train a recurrent network that reproduced natural speech and generated both slower and faster utterances of natural words with no additional training. Using further refinements of the model, we employed this principle of oscillation-driven network to capture the fast replay of place cells during a task of spatial navigation.

Below we discuss the biological implications of our model as well as its applications and limitations.

### B. Biological relevance and predictions of the model

Despite some fundamental limitations common to most computational models of brain activity, our approach was designed with several key features of living neuronal networks, including spiking neurons, Dale’s principle, balanced excitation/inhibition, a heterogeneity of neuronal and synaptic parameters, propagation delays, and conductance-based synapses (7, 11, 46). We used a learning algorithm that isn’t biologically plausible to train the readout unit (recursive least-square). However, given that the SRNN’s dynamics is independent of the readout’s output, any other learning algorithm would be compatible with our model.

Further, and most central to this work, our model included neural oscillations along a range of frequencies that closely matched those reported in electrophysiological studies (30, 47). Although there is an abundance of potential roles for neural oscillations in neuronal processing, much of their function remain unknown (48). Here, we proposed that multiple heterogeneous oscillations may be combined to generate an input whose duration greatly exceeds the time-course of any individual oscillation. In turn, this multiplexed input allows a large recurrent network operating in the chaotic regime to generate repeatable and stable patterns of activity that can be read out by downstream units.

It is well established that central pattern generators in lower brain regions such as the brain stem and the spinal cord are heavily involved in the generation of rhythmic movements that match the period of simpler motor actions (e.g. walking or swimming) (49). From an evolutionary perspective, it is compelling that higher brain centers would recycle the same mechanisms (47) to generate more complex and non-repetitive actions (50, 51). In this vein, (51) showed that both periodic and quasi-periodic activity underlie a non-periodic motor task of reaching. Our model provides a framework to explain how such activity can be exploited by living neuronal networks to produce rich dynamics whose goal is to execute autonomous aperiodic tasks.

While our model shows that oscillatory networks can generate input oscillations that control the activity of a SRNN (Fig. 2), the biological identity of these oscillatory networks is largely circuit-dependent and may originate from either intrinsic or extrinsic sources. In the hippocampus, computational (39) and experimental (40) findings suggest an intrinsic source to theta oscillations. Specifically, studies raise the possible role of CA3 in forming a multi-periodic drive consisting of several interdependent theta generators that activate during spatial navigation (41).

Similarly, the neural origin of the tonic inputs controlling the activity of the oscillatory networks is not explicitly accounted for in our simulations. However, it is well established that populations of neurons can exhibit bistable activity with UP-states lasting for several seconds (52) that could provide the necessary input to drive the transition from asynchronous to synchronous activity in oscillatory networks.

Going beyond an *in silico* replication of neurophysiological findings, our model makes two empirically testable predictions. If one was to experimentally isolate the activity of the input oscillators, one could show that: (*i*) a key neural signature of a recurrent circuit driven by multi-periodic oscillations is the presence of inter-trial correlations between the phase of these oscillations; and (*ii*) the period of the input oscillators should appear faster or slower to match the rescaling factor of the network. This correspondence between the input oscillations and temporal rescaling is a generic mechanism behind the model’s ability to perform a wide variety of tasks, from spatial navigation to speech production.

### C. Related models

Our approach was inspired by predecessors in computational neuroscience. Multiplexing multiple oscillations as a way to generate long sequences of non-repeating inputs was first introduced by (18), with a model of interval timing relying on the coincident activation of multiple oscillators of different frequencies. This idea served as a basis for the striatal beat frequency model (19), where multiple cortical regions are hypothesized to project to the striatum which acts as a coincidence detector that encodes timing intervals. A similar mechanism was also suggested for the representation of space by grid cells in rodents (17). Grid cells have periodic activation curves spanning different spatial periods (53), and their activation may generate a combinatorial code employed by downstream regions to precisely encode the location of the animal in space (17).

Other studies have suggested that phase precession during spatial navigation could originate from a dual oscillator process (43, 54). Along this line, a recent model of the hippocampus uses the interference between two oscillators to model the neural dynamics related to spatial navigation (55). Although this model shares similarities with ours, a fundamental difference is that our model uses the phase of combined oscillators to create a unique input at every time-step of a task, whereas their model relies on the beat of the combined frequencies. Additionally, in our model, increasing the frequency of input oscillations by a common factor leads to compressed sequences of activity. By comparison, in the model of (55), sequences are compressed by removing an extrinsic input oscillator. More experimental data will be needed to support either model.

### D. Limitations and future directions

In our model, periodic activity was readily observable in the SRNN dynamics due to the input drive (e.g. Fig. 1c or Fig. 6d). However, the architecture of our model represents a simplification of biological networks where several intermediate stages of information processing occur between sensory input and behavioral output. Oscillatory activity resulting from a multi-periodic drive might occur in one, but not necessarily all stages of processing. Further work could examine this issue by stacking SRNN connected in a feed-forward manner; such a hierarchical organization may have important computational benefits (56).

In the spatial navigation task, we ensured that the location of the animal was perfectly correlated with the time spent in the place field of each cell. This is, of course, an idealized scenario that does not account for free exploration and variable speed of navigation along a track. These factors would decorrelate the spatial location of the animal and the time elapsed in the place field. Hence, further work would benefit from a more ecologically-relevant version of the navigation task. This new version of the task might aim to capture how the time spent in a given place field impacts the link between the activity of place cells and theta oscillations (57).

Finally, our task of speech production was restricted to learning a spectrogram of the target signal. This simplified task did not account for the neural control of articulatory speech kinetics, likely involving the ventral sensory-motor cortex (58, 59).

#### D.1. Applications

Our modelling framework is poised to address a broad spectrum of applications in machine learning of natural and artificial signals. With recent advances in reservoir computing (60) and its physical implementations (61), our approach offers an alternative to using external arbitrary time-varying signals to control the dynamics of a recurrent network. Our model may also be extended to neuromorphic hardware, where it may benefit chaotic networks employed in robotic motor control (62). Finally, our model is, to our knowledge, the first to produce temporal rescaling of natural speech, with implications extending to conversational agents, brain-computer interfaces, and speech synthesis.

Overall, our model offers a compelling theory for the role of neural oscillations in temporal processing. Support from additional experimental evidence could impact our understanding of how brain circuits generate long sequences of activity that shape both cognitive processing and behavior.

### E. Data availability

The custom python scripts used to replicate the main results of this manuscript can be found at https://github.com/lamvin/Oscillation_multiplexing.git.

## ACKNOWLEDGMENTS

This work was supported by a doctoral scholarship from the Natural Sciences and Engineering Research Council (NSERC) to P.V.L., the program of scholarships of the 2nd cycle from the Fonds de recherche du Québec – Nature et technologies (FRQNT) to M.C., and a Discovery grant to J.P.T. from NSERC (Grant No. 210977).

## Supporting Information

### Parameter exploration of the SRNN

Fig.S1 shows the impact of different network parameters on the model’s performance. We used three measures of the network’s activity and performance: 1-The average coefficient of variation (CV) between trials for each cell. A low CV shows that the ratio of across-trial variability is low relative to the mean firing rate. 2-The average firing rate across all trials. 3-The average correlation between the output of the network and the target function after training.

### Autonomous production of oscillatory activity

For a given network, it is possible to modulate the frequency of its activity by tuning the gain of the tonic input it receives. Stronger external inputs lead to a faster period, up to a limit after which the stability of the activity degrades (Fig.S2a,b). The activity of oscillatory networks remains stable in the presence of noise, as the network’s cells received trains of synaptic inputs drawn from a Poisson distribution with increasing frequencies and the networks were all robust to clamping a subset of their neurons to their resting potential for the duration of the simulation (Fig.S2c).

We explored the impact of network parameters on the production of repeatable periodic activity. We were interested in networks that were spontaneously active and that switched to a synchronized regime when driven by a constant tonic input. Starting with networks identical to the main SRNN except for its size (500 cells instead of 1000), we tuned the gain and time-scale of inhibitory inputs as well as the gain of the external input and the network sparseness, and monitored the dynamics of the network. We found that all those parameters were crucial in shaping the transition between oscillatory and asynchronous regimes.

To assess to repeatability of network activity, we computed the Pearson correlation of the firing rate fluctuations during each trial with the first trial for every network (Fig.S3). Overall, regions of lower inhibition conductance and higher inhibitory synaptic time-constants favour the emergence of repeatable oscillations. The strength of the tonic input also had an impact on the network activity.

### Rescaling slopes

We examined whether the qualitative differences in performance slopes between faster and slower rescaling is reflected in the SRNN’s activity. We first obtained the projection of the SRNN’s activity on the first principal component (accounting for most of the variance of the SRNN, which is typical for this type of network (63)). We then scaled back the activity of the first principal component to the original scaling for all the rescaling factors tested (Fig.S6a). This allowed us to compare the features of the SRNN’s activity at the original scale. We used a Fourier analysis to obtain a representation of the SRNN’s activity for all rescaling factors at different frequencies (Fig.S6b). For each rescaling and each frequency band tested, we computed an “error” value defined as the mean squared difference between the SRNN’s activity scaled back to the original velocity and the activity of the SRNN on the original scaling at a given frequency. Each error value was then normalized by the square of the frequency band for the target (thus making comparisons of error across frequencies possible).

This analysis revealed that the higher frequencies of the network activity deteriorated faster than the lower frequencies. The lower frequencies were well-preserved for inputs that were slowed down, but were progressively lost for faster rescalings. This explains why the performance reached a plateau (in the absence of input noise) for longer rescaling. We used the same normalization process to examine the output performance across different velocities (Fig.S6c). We computed the error for each frequency until 6 Hz given that the target function was generated by applying a low-pass filter on white noise at this cut-off. All frequencies degraded evenly for faster velocities. However, unlike the SRNN’s activity, the higher frequencies appeared to be more preserved for the output, whereas the lower frequencies (<4Hz) were more degraded. These results show that rescaling the input frequencies lead to complex modifications of the SRNN dynamics, and highlight a non-trivial relationship between the input and output of the model.

### Input oscillations and phase jitter

We simulated a random-walk process *ϕ*(*t*) (see Methods) with a standard deviation *σ*_*ϕ*_ that we added to the phase of an oscillator (Fig.S7a). Because the noise of the random walk comes from a Gaussian process with mean 0, the expected deviation from the base sinusoid is null at all time steps, but its standard deviation at time *t* is equal to 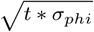 (Fig.S7b).

We then trained our model with different *σ*_*phi*_ values for different durations. Fig.S7c shows the expected standard deviation in degrees for the *σ*_*ϕ*_ used in our simulation, and well as the corresponding error between the output and target functions. Our results show that error accumulates exponentially (*R*^2^ = 0.92, Fig.S7d). This means that networks can tolerate a deviation of up to 50 degrees from the original sine wave before reaching random performance (defined as the plateau in MSE of the output in Fig.S7c,d).

**Fig. S1.**
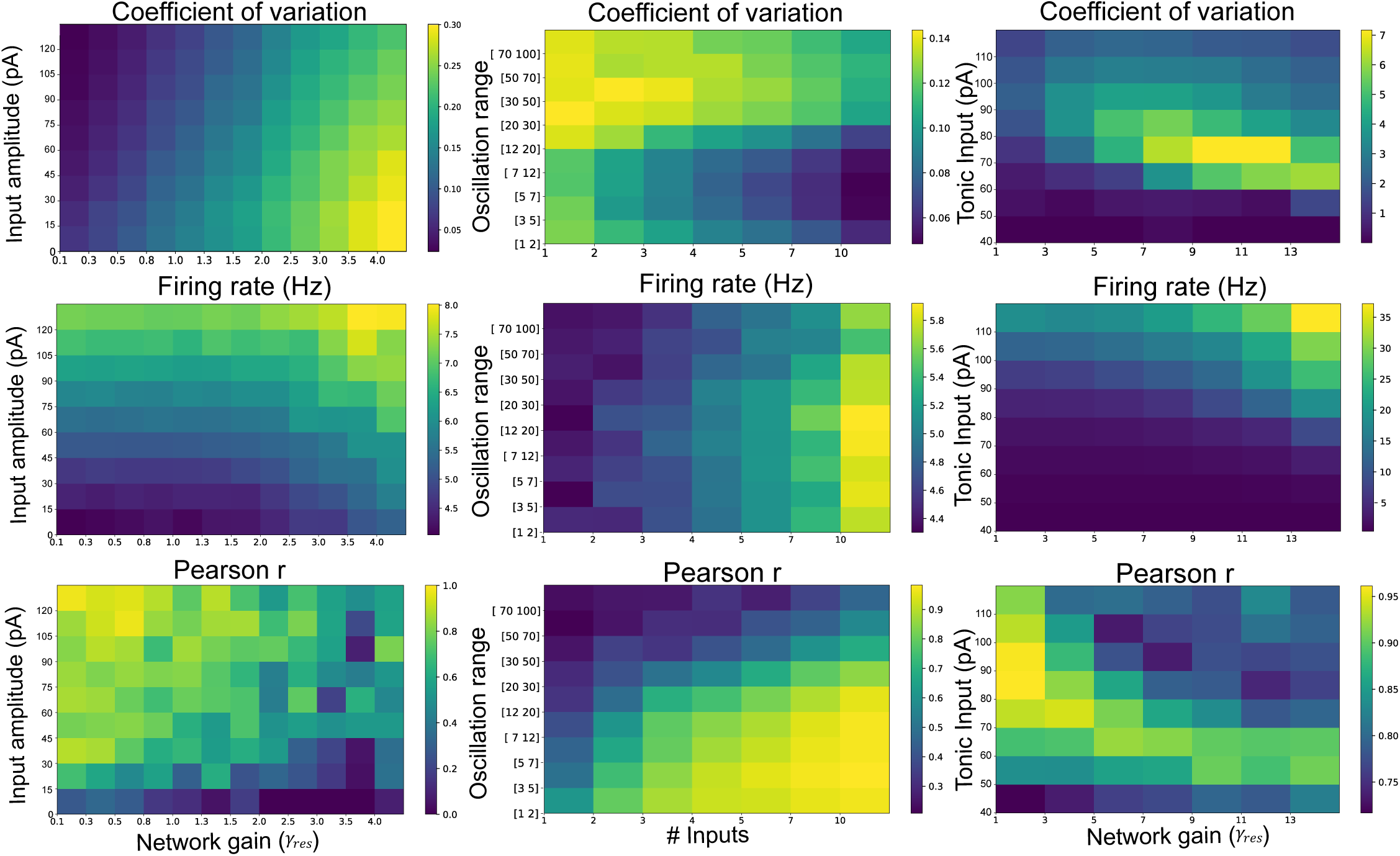
Parameter exploration for training a SRNN. The top row shows the average CV, the middle row shows the average firing rate, and the bottom row shows the average correlation between the output of the network and the target output. *Input amplitude* is the strength of the the input projecting to the SRNN’s units. *Network gain* is the strength of the recurrent connections between the SRNN’s units. *Oscillation range* is the range from which each input unit’s sine wave period is randomly drawn. *# Inputs* is the number of different input units projecting to the SRNN. *Tonic input* is the strength of the constant current injected to each of the SRNN’s units.

**Fig. S2.**
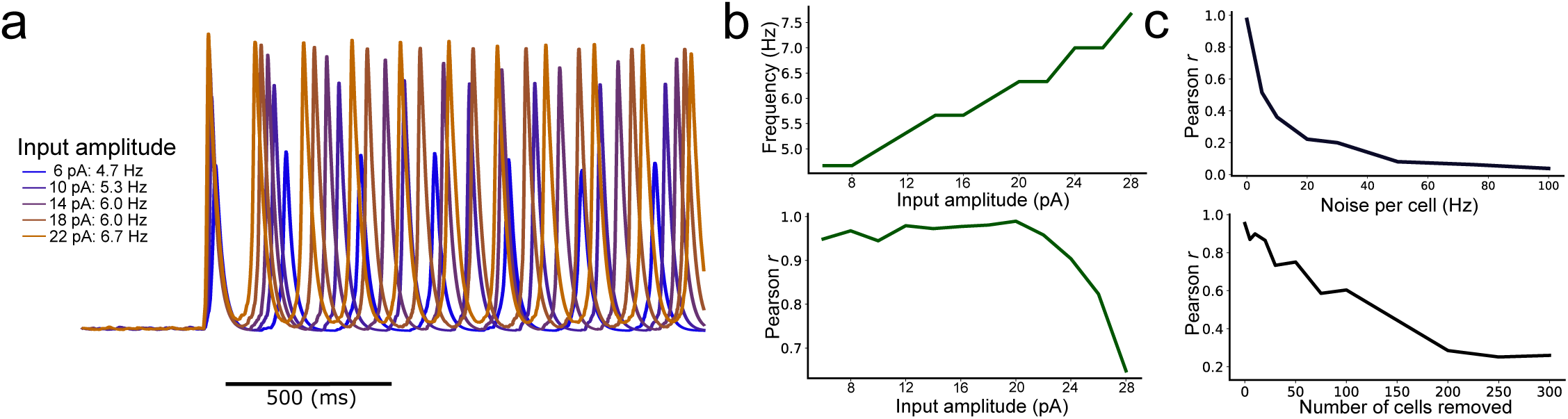
Autonomous production of repeatable periodic activity. **a** Average current over all neurons for five trials of each condition as a function of the strength of the external drive. The frequency scales with the strength of the tonic input. **b** Top: Average frequency of the network’s activity as a function of the external drive. Bottom: Pearson correlation between multiple trials as a function of the amplitude of the external drive. **c** Pearson correlation between multiple trials for a given network as a function of the rate of noisy synaptic input per neuron (top) and as a function of the proportion of clamped neurons (bottom).

**Fig. S3.**
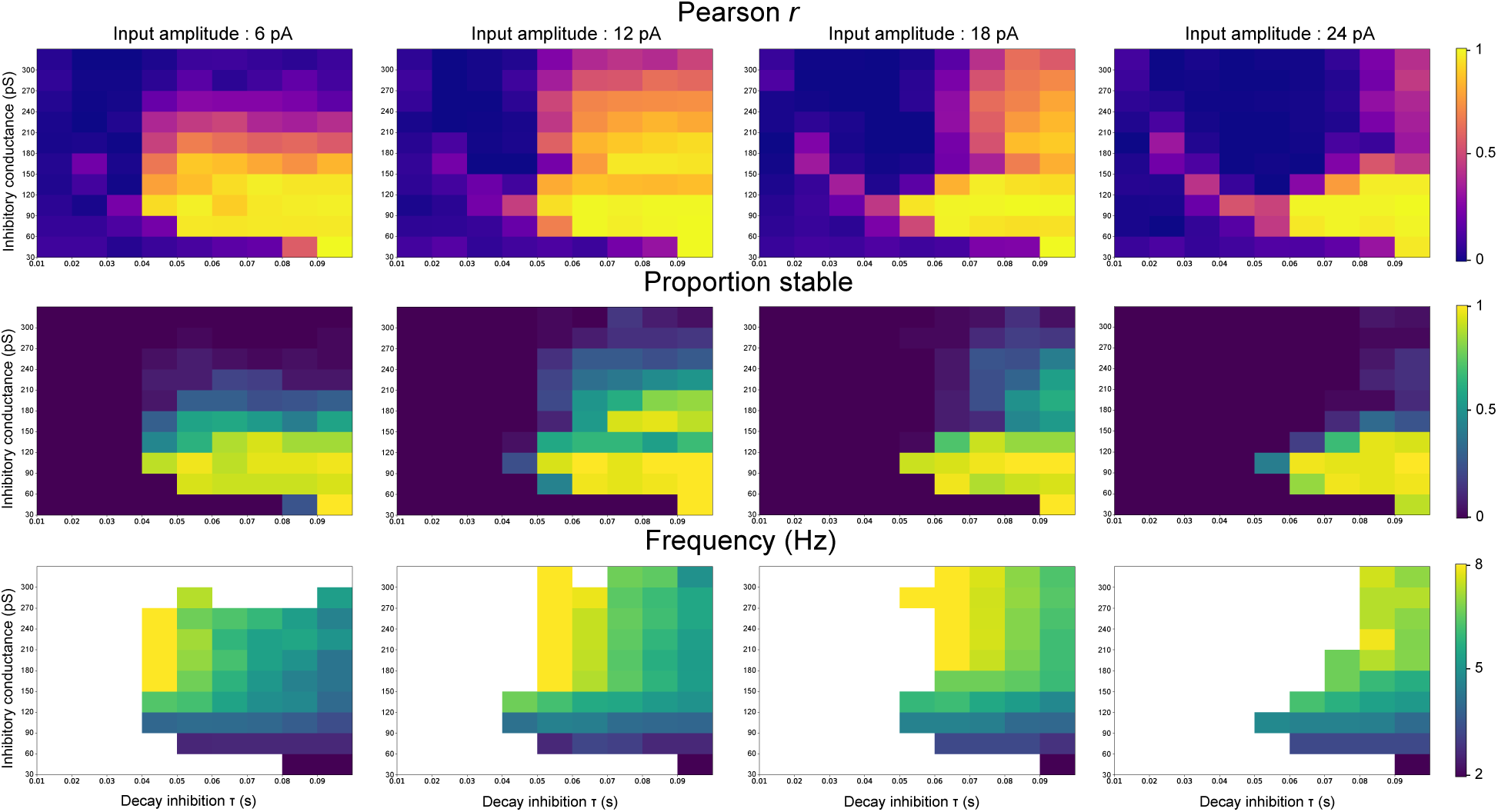
Parameter exploration of the network’s synchronous activity (inhibition conductance and kinetics). Each column represents a different strength of the external input step, and each heatmap shows different combinations of the conductance and synaptic time-constant of inhibitory inputs. The top row shows the average Pearson correlation between the output and the target function for 5 trials for each of the 50 random network initializations tested per condition. The middle row shows which proportion of the networks are considered as stable. The bottom row shows the average frequency of the periodic activity of the stable networks.

**Fig. S4.**
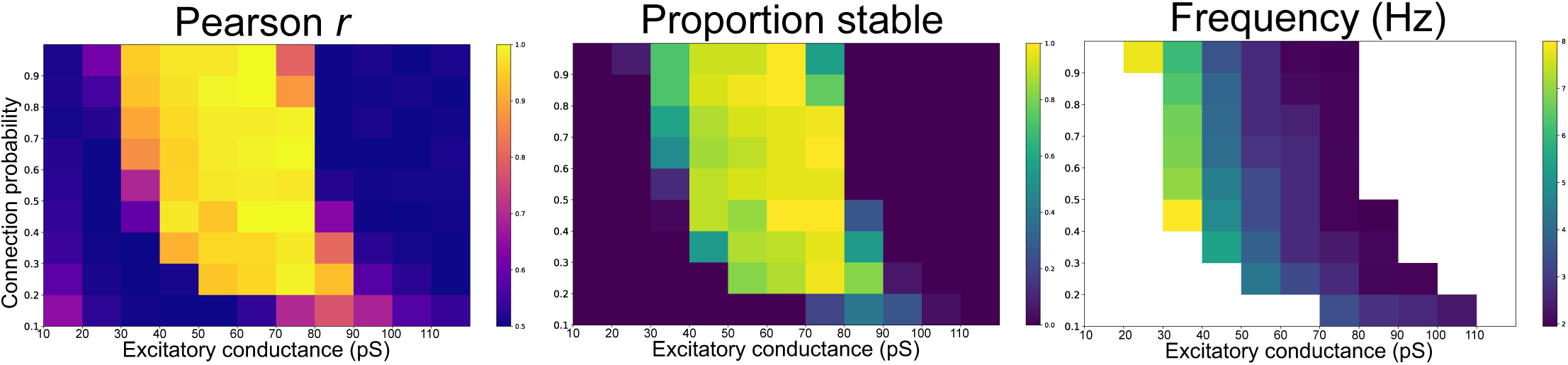
Parameter exploration of synchronous activity (excitatory conductance and *p*). The sparseness of the SRNN and the conductance of the excitatory connections were systematically varied to observe their impact on the network’s regime. *Left* : Pearson correlation between the output and the target function for 5 trials for each of the 50 random network initializations tested per condition. *Middle*: proportion of networks that are considered stable. *Right* : average frequency of the periodic activity of the stable networks.

**Fig. S5.**
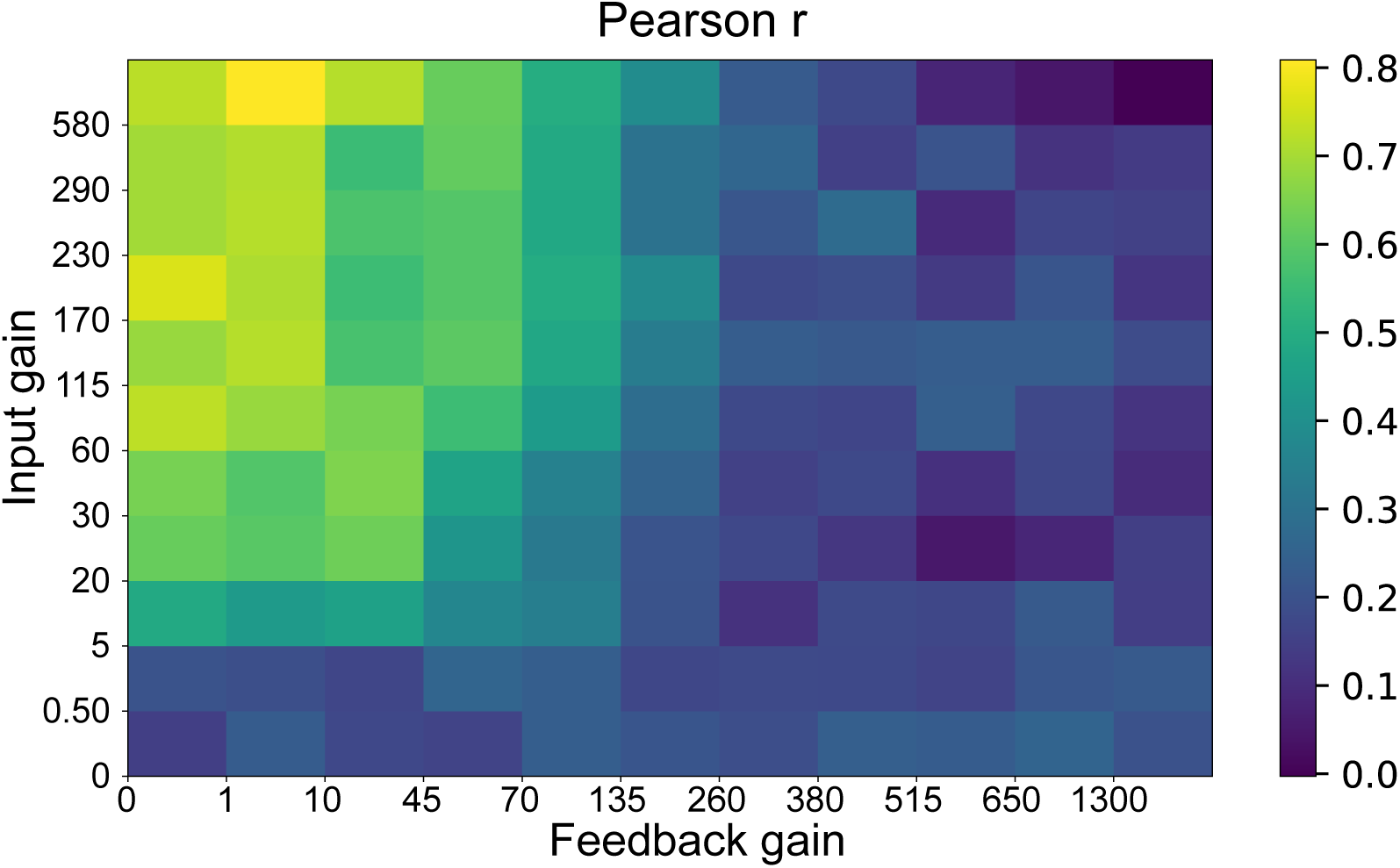
Impact of SRNN feedback to the oscillatory networks. The models can perform well when the gain of the feedback connections from the SRNN to the oscillatory networks are scaled up to a limit. The effect of the feedback gain appears to be independent of the strength of the tonic input of the step function to the oscillatory networks.

**Fig. S6.**
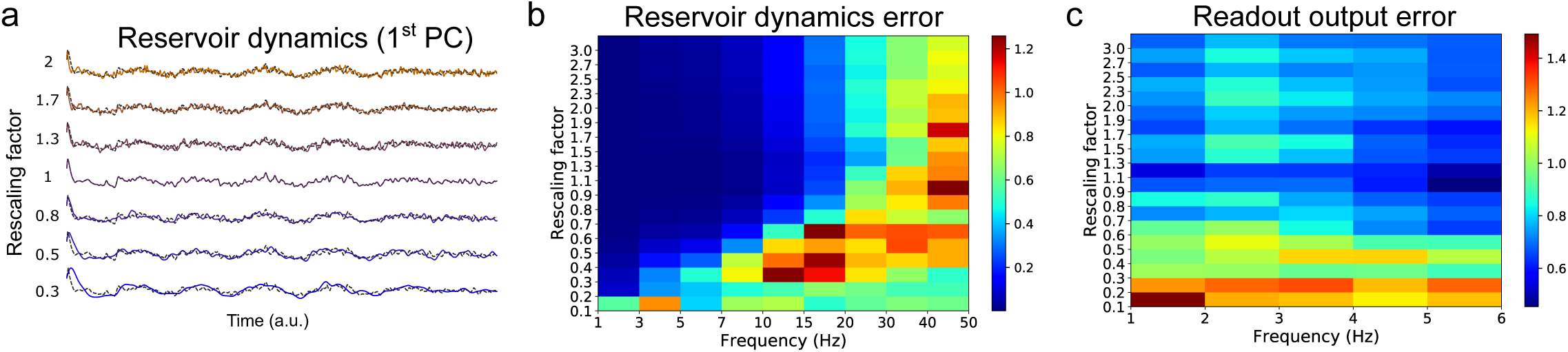
Network dynamics and performance with temporal rescaling. **a** Activity of the SRNN projected on the first principal component for different rescaling factors. The black dashed line represents the activity without rescaling. The traces displayed have been scaled back to the original velocity. **b** Normalised error of the first principal component of the SRNN’s activity for different rescaling factors and frequency bands. **c** Normalised error of the network’s output (MSE between the output and the target divided by the total variance of the target function) for different rescaling factors and frequency bands.

**Fig. S7.**
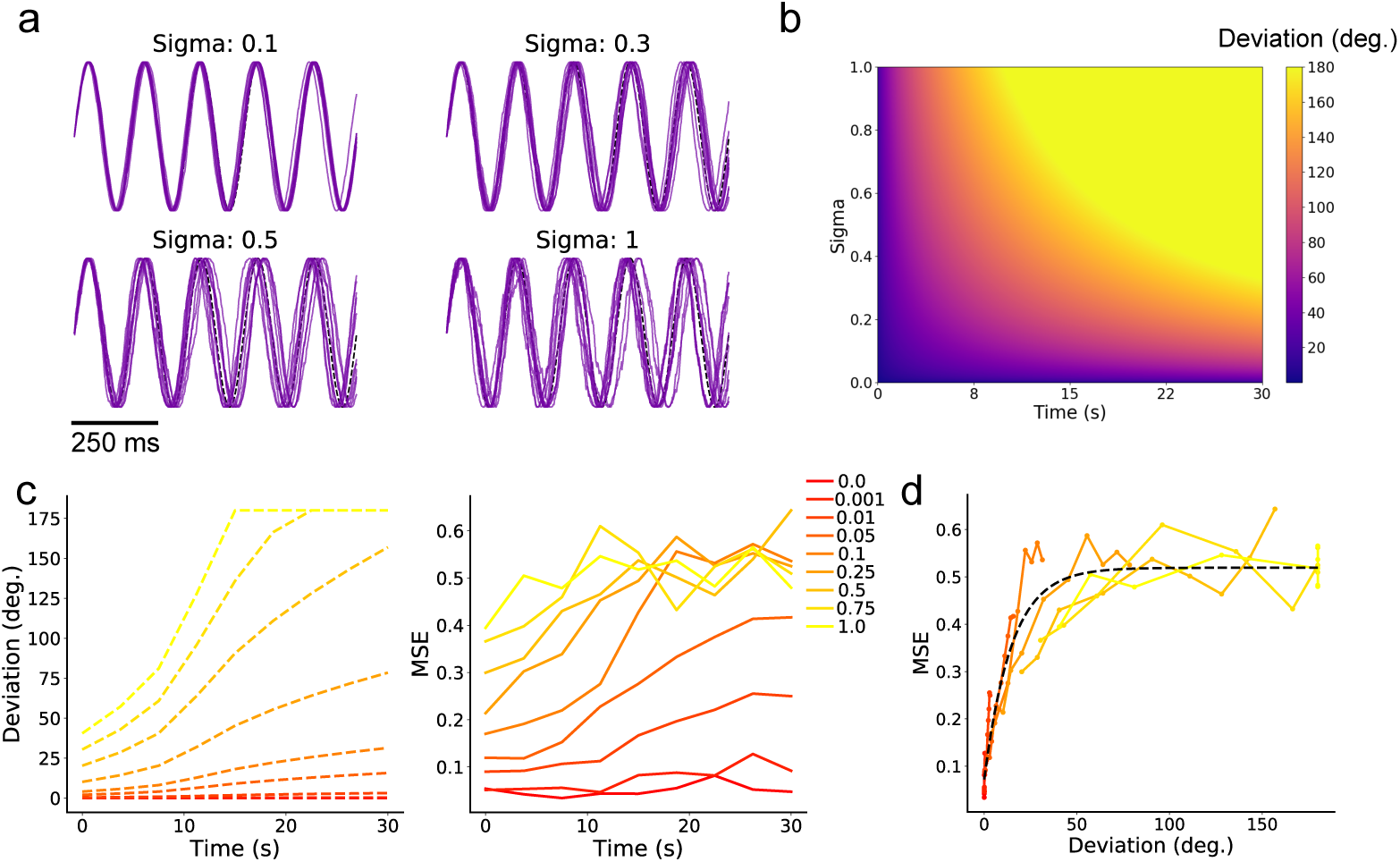
Network performance with desynchronized inputs. **a** Four simulations of 10 trials each with different *σ*_*ϕ*_ with a random walk process added to the phase of a sine wave. **b** Analytical standard deviation (in degrees) around the expected phase of a deterministic signal as a function of time at different values of *σ*_*ϕ*_. **c** Left: Analytical estimation of the standard deviation in degrees with the different *σ*_*ϕ*_ used for the network simulation. Right: corresponding MSE between the output and target of the readout unit for the different conditions tested. **d** Relationship between the deviation in degree and the MSE. An exponential function was fit to the data (*R*^2^=0.92).

